# Comparative Population Genetics in the Human Gut Microbiome

**DOI:** 10.1101/2021.03.02.433642

**Authors:** William R. Shoemaker, Daisy Chen, Nandita R. Garud

## Abstract

The genetic variation in the human gut microbiome is responsible for conferring a number of crucial phenotypes like the ability to digest food and metabolize drugs. Yet, our understanding of how this variation arises and is maintained remains relatively poor. Thus, the microbiome remains a largely untapped resource, as the large number of co-existing species in this microbiome presents a unique opportunity to compare and contrast evolutionary processes across species to identify universal trends and deviations. Here we outline features of the human gut microbiome that, while not unique in isolation, as an assemblage make it a system with unparalleled potential for comparative population genomics studies. We consciously take a broad view of comparative population genetics, emphasizing how sampling a large number of species allows researchers to identify universal evolutionary dynamics in addition to new genes, which can then be leveraged to identify exceptional species that deviate from general patterns. To highlight the potential power of comparative population genetics in the microbiome, we re-analyzed patterns of purifying selection across ~40 prevalent species in the human gut microbiome to identify intriguing trends which highlight functional categories in the microbiome that may be under more or less constraint.

## Introduction

The human microbiome is a complex ecosystem composed of hundreds of interacting species. Although the diversity of the microbiome has been extensively studied at a species level, each species harbors genetic diversity that is quite varied across hosts as well as within a host over time (Zhu et al. 2010, 2019; Schloissnig et al. 2013; Faith et al. 2013). This genetic diversity can confer a number of crucial traits to microbes as well as their hosts, such as the ability to digest food, metabolize drugs, and evade antibiotics. However, our understanding of how these genetic variants arise and segregate via population genetic forces – e.g., random genetic drift, mutation, recombination, selection, and migration – across the hundreds of species that call our guts home, is relatively nascent (Garud & Pollard 2020).

Our knowledge of how evolution proceeds in a community context is similarly underdeveloped. Much of our intuition about the evolution of microorganisms come from studying individual species (Good et al. 2017; Bruger & Marx 2018; Herron & Doebeli 2013; Tenaillon et al. 2016; Xue et al. 2017; Lieberman et al. 2014). By contrast, the microbiome is composed of hundreds of interacting species and strains in which both ecological and evolutionary forces simultaneously act (Garud & Pollard 2020; Good & Hallatschek 2018). Specifically, change in the frequency of an existing haplotype as well as the emergence of a new haplotypes (i.e., evolution) can occur on the same timescale as changes in strain frequencies (i.e., ecology). These simultaneous ecological and evolutionary processes within the human gut microbiome affords a unique opportunity. Rather than studying one species at a time, we can study the population genetics and ecology of many (≳40) species almost simultaneously. Thus, the human gut microbiome is a model system for studying comparative population genetics across co-existing species in a natural environment.

Comparative population genetics is a rich area of study that has yielded insights into novel functional elements and population genetic processes in macroscopic organisms ranging from mammals (Romiguier et al. 2014; Lindblad-Toh et al. 2011; Pollard et al. 2010; Davydov et al. 2010) to *Drosophila melanogaster* (Clark et al. 2007; Lawrie & Petrov 2014). Now with the availability of hundreds of thousands of new genomes from the deep sequencing of commensal microbes (Pasolli et al. 2019; Almeida et al. 2019; Nayfach et al. 2019), similar comparative analyses can be made across microbial species and populations. However, unlike typical macroorganisms, the microbiome provides a rich opportunity to understand how ecological interactions between species modulate evolutionary dynamics within individual species.

### A model system for comparative population genetics: the human gut microbiome

The human gut microbiome is a compelling system for comparative population genetics. The short time scales on which evolutionary dynamics occur in the microbiome make it possible to witness evolution in action as well as how evolution interacts with ecology (Zhao et al. 2019; Garud et al. 2019; Yaffe & Relman 2020; Poyet et al. 2019; Roodgar et al. 2020). Moreover, the meta-population structure of human microbiomes lends itself naturally to treating measurements in spatially distant hosts as independent biological replicates, from which we can decode general principles. Given the massive amounts of data now available from thousands of individuals from around the world (Almeida et al. 2019; Pasolli et al. 2019), the time is ripe for the study of evolution in the microbiome. Additionally, the ability to experimentally manipulate this system (Roodgar et al. 2020) as well as its members (Barroso-Batista et al. 2014; Zhao et al. 2019; Ramiro et al. 2020) enables validation of computational predictions and discovery of new principles. Finally, the strong relevance of the microbiome to our health makes it a medically important system (Thomas et al. 2019; Ley et al. 2005; Jakobsson et al. 2010; Duvallet et al. 2017). Thus, comparative population genetic studies in the microbiome will not only elucidate general principles relevant to the microbiome, but also the broader field of population genetics. Here we elaborate on these fundamental facets of the microbiome that make it the ideal system to study comparative population genetics in a naturally complex ecosystem.

#### 1) Rapid evolution on short timescales

With relatively short generation times of only(~1-10 days in the human gut (Korem et al. 2015), microbes have the potential to rapidly evolve. This enables temporally resolved analyses that are almost impossible to replicate in charismatic macroscopic organisms. A sense of the evolutionary timescale of the human microbiome is best illustrated through a back-of-the-envelope calculation: taking the low end of the range of generation times for the gut microbiome of ~1 day (Korem et al. 2015; Milo & Phillips 2016) we find that it has evolved for ~9,000 generations before its host has reached age 25, a number that is on the order of the ~10,000 generations that humans have existed (Moorjani et al. 2016; Scerri et al. 2018). Alternatively stated, the microbiome within a human host can evolve on a timescale similar to that of the entire human species.

The ability to sample populations over a large number of generations can alter how evolutionary biologists approach their questions. Instead of relying predominantly on phylogenetic reconstruction from static data, researchers can effectively observe evolution in real time. Over the course of just a few months, new genotypes can emerge and recombine over short time scales, and ultimately become lost or fixed over extended time scales. Indeed, it recently has been observed that adaptation and recombination can occur in the human gut over a matter of months (Zheng et al. 2020; Yaffe & Relman 2020; Poyet et al. 2019; Garud et al. 2019; Zhao et al. 2019; Lin & Kussell 2017). However, the necessary temporal resolution of sampling remains subject to the researcher’s question, as short-term adaptation in response to a temporary environmental perturbation (e.g., a host consuming antibiotics over a few weeks) may require denser sampling than evolution in a relatively unchanging environment.

#### 2) Replication across hosts

Given that the human gut microbiome is a quasi-closed system, separate hosts can be treated as replicate evolution studies, an observation that has been summarized by the captivating moniker “seven billion microcosms” (Lieberman 2018). This level of replication can be leveraged to identify targets of positive selection that recurrently accumulate fixation events in independent hosts (i.e., parallel evolution; Xue et al. 2017; Bertels et al. 2019; Lieberman et al. 2014; Zhao et al. 2019; Poyet et al. 2019). By combining large cohort sizes with temporally resolved sampling we can also examine how these signatures of parallelism change over time, allowing us to dissect the temporal dynamics of adaptation (Barroso-Batista et al. 2014). For example, targets of rapid adaptation typically harbor a disproportionate number of sites with strong beneficial fitness effects, leading to the fixation of multiple mutations within a short period of time (i.e., "coupon collecting"; Good et al. 2017). Alternatively, if the fitness effects of sites in a gene depend on whether prior mutations have fixed elsewhere in the genome, then the time between fixation events will be large (i.e., historical contingency; Gould 1990; Blount et al. 2008). The benefits of large cohorts are not limited to detecting adaptation. With large cohorts, we can also observe deleterious alleles segregating at extremely low frequencies that are likely subject to purifyin selection (Lawrie & Petrov 2014). Combined with exciting recent theoretical developments (Neher & Shraiman 2012; Nicolaisen & Desai 2012; Cvijović et al. 2018; Good 2020), the human gut microbiome is an ideal system to examine the evolutionary dynamics of purifying selection.

#### 3) Ecology and evolution frequently interact

It is becoming increasingly clear that mutations in many microbial populations do not fix or become extinct, instead remaining at intermediate frequencies for extended periods of time (Good et al. 2017; Good & Hallatschek 2018). This “strain” level structure constitutes a form of ecology that exists below the taxonomic level of species, and is commonly found within hosts for most gut microbiota (Garud et al. 2019). The sheer prevalence of strain structure in the human microbiome and the fact that they can differ on the order of a few nucleotides (Goyal et al. 2021) suggests that ecological and evolutionary dynamics occur on similar timescales in microbial systems, contrary to the historical belief that evolutionary timescales are longer than ecological timescales (Slobodkin 1980). For example, strain frequencies can fluctuate on the same time scale on which they acquire new genetic adaptations (Garud et al. 2019; Zhao et al. 2019). This observation, along with the relative ease with which a large number of species can be sampled across hosts, suggests that the human gut microbiome is a system with unmatched potential for the exploration of eco-evolutionary dynamics.

The presence of overlapping ecological and evolutionary timescales in the microbiome has spurred empirical and theoretical efforts to characterize eco-evolutionary interactions within the gut. A prime example being a recent mathematical model that describes how the frequency of a strain can change over time as *de novo* mutations accumulate, which affect how said strain consumes environmentally supplied resources in addition to its overall fitness (Good et al. 2018). Thus, evolution can affect competition between strains with resource consumption as a mediating factor, changes that ultimately alter community composition and structure. However, it is unlikely that eco-evolutionary interactions within the gut can be sufficiently captured by accounting for environmentally supplied resources alone. Rather, microorganisms often secrete secondary metabolic compounds, supplying additional resources that can be consumed by other species (i.e., cross-feeding). This metabolic dependency promotes species co-existence and becomes increasingly likely in communities with many species, an ecological bedrock that supports subsequent coevolution (D’Souza et al. 2018; Lilja & Johnson 2019). The widespread nature of this phenomenon may explain empirical patterns where it appears as though the arrow of causation between evolution and ecology is reversed, a prominent example being that the diversification rate of a species is correlated with the number of species in its community (Madi et al. 2020).

#### 4) Experimental manipulation

While a natural system enables researchers to study complex phenomena that cannot be recapitulated exactly in the laboratory, some degree of experimental manipulation is necessary to validate predictions and generate new insights. Over the last few years, substantial progress has been made towards characterizing the evolutionary dynamics of adaptation in microbial populations. The use of lineage tracing via barcoding has allowed the distribution of fitness effects of *de novo* mutations to be quantified in certain species (Levy et al. 2015) in addition to providing evidence that the travelling wave is an appropriate model of microbial adaptation (Nguyen Ba et al. 2019). The ease with which such techniques can be applied to non-model species varies, though gene deletion via transposon mutagenesis libraries has been shown to be particularly effective for identifying loci that confer a growth advantage in environments with different resources (Cain et al. 2020), different sets of co-occurring species (Thibault et al. 2019), for species isolated from the gut (Ruiz et al. 2013), and in the gut microbiome of model organisms (Powell et al. 2016; Zimmermann et al. 2019; Ludington & Ja 2020; Barreto et al. 2020; Barroso-Batista et al. 2020). These experiments can serve as a compliment to traditional comparative population genetic analyses, allowing us to test hypotheses formed from metagenomic observational studies. While all these studies fall short of true *in vivo* manipulation of human guts, they are useful approximations that allow for high-throughput manipulations to be performed and evolutionary and ecological hypotheses to be tested.

#### 5) Relevance to health

A significant benefit to studying comparative population genetics in the human gut microbiome is that the findings made may have a direct relevance for human health. The species composition of the gut microbiome is known to be essential for proper immunological (Belkaid & Hand 2014), neurological (Yano et al. 2015), and metabolic development (Rowland et al. 2018), and is associated with a number of human diseases including colorectal cancer, diabetes (Vallianou et al. 2018), and obesity (Ley et al. 2005, 2006). While the connection of the human microbiome to host health has been primarily studied at the species level, genetic variants in the microbiome play a crucial role for health as well. Specifically, microbiome genetic variants can confer a number of critical traits to human hosts, including the digestion of new foods (Kenny et al. 2020; Hehemann et al. 2010), antibiotic resistance (Gautam et al. 2018), and the metabolization of drugs (Spanogiannopoulos et al. 2016). A comparative genomics approach will enable the discovery of new microbiome genetic variants (Sberro et al. 2019), which may ultimately be useful for the future development of effective microbiome therapies.

### Lessons from comparative population genetics in the microbiome

While our understanding of the microbiome and the discipline of comparative population genetics have rapidly expanded since the emergence of next-generation sequencing almost two decades ago, their intersection is relatively recent. Therefore, the potential of comparative population genetics in the microbiome is still being realized. Here we present three goals for the future of comparative population genetics: the need to identify 1) previously unknown functional elements of microbial genomes, 2) evolutionary dynamics common to all species in the gut microbiome, and 3) individual species and genomic features that deviate from general patterns.

#### 1) Inference of Functionality

Currently, the annotation of genes in the microbiome and our understanding of their functionality is severely lacking, with an estimated 40% being “hypothetical” (Almeida et al. 2019). Comparative population genetics has the potential to shed light on the functions of existing hypothetical genes and assist with the identification of new ones. Much of the utility of comparative population genetics derives from the neutral theory of molecular evolution, which predicts that if mutations in functional regions of genomes tend to be deleterious, those regions will evolve at a slower rate than effectively neutral nonfunctional regions (Kimura 1983). This constraint allows for conserved elements of the genome to be identified between highly diverged species; a “phylogenetic footprint” (Lawrie & Petrov 2014). Using this basic assumption, comparative genomic analyses across groups of macroorganisms as diverse as *Drosophila* and mammals have yielded insight into novel proteins (Clark et al. 2007; Lawrie & Petrov 2014). Now, recent efforts have been made to apply this approach to the microbiome (Fremin & Bhatt 2020). Specifically, Sberro et al. (2019) recently performed a comparative analysis on shotgun microbiome metagenomic data and discovered thousands of novel small genes. Among their discoveries was a novel small ribosome-associated protein that seems to be transcribed and translated at high levels. Despite the fundamental functional significance of this protein, it may have been missed due to the historical focus on model organisms such as *E. coli* and common pathogens.

However, there is additional justification to claim that microorganisms harbor a substantial number of unannotated functional elements. Population genetic theory coupled with cellular energetics predicts that the vast majority of unannotated genes within the gut microbiome likely play some functional role (Lynch & Marinov 2015; Martinez-Gutierrez & Aylward 2019). For example, a single nonfunctional nucleotide within a microbial genome is visible to purifying selection (Lynch & Marinov 2015), a stark contrast to macroorganisms where junk DNA is highly prevalent (Lynch 2007). Coupled with the higher gene density in microorganisms due to overlapping open reading frames (Johnson & Chisholm 2004), this prediction suggests that the gut microbiome is a particularly apt system for researchers who wish to leverage statistical evidence provided by comparative population genomics to confirm the purported functionality of a given gene. Indeed, researchers are likely already acting on this prediction, as recent efforts combined comparative genomics with RNA-seq to identify ~2,000 novel structural RNAs in the microbiome (Fremin & Bhatt 2020). With hundreds of species harboring genomes with high gene density across billions of hosts, the gut microbiome is still very much a proverbial “gold rush” for the discovery and characterization of novel proteins and RNAs.

#### 2) Robust evolutionary patterns

Arguably, it is necessary to gain some degree of knowledge regarding the typical evolutionary dynamics of a species in a given system before comparisons between species can be performed. At first glance, it would appear as if the goal of identifying general evolutionary patterns in the human gut microbiome is hopeless. There are few cases where population geneticists would say that we have sufficient knowledge of the evolutionary dynamics of one species, much less hundreds or thousands of species that interact in the same environment. Operating under this assumption, we would conclude that the complexity of the microbiome is irreducible. This is not entirely an unjustified claim, since if one is interested in the evolutionary dynamics of an individual species, how can those dynamics be sufficiently characterized if you cannot examine an isolated species *in vivo*?

The fault here is the idea that we need to understand the dynamics of individual species to understand the general dynamics of the system. Instead, progress can be made by temporarily abandoning the Cartesian framework that is ubiquitous in traditional biology (Levins & Lewontin 1987) and embracing an alternative approach, where we exchange determinism for a statistical property, the average over an ensemble of species. This rationale is essentially what physicists realized in the 19^th^ century (Pathria & Beale 2011) and has been applied in recent years to examine the ecological dynamics of microorganisms, through the development of mathematical models (Advani et al. 2018; Barbier & Arnoldi 2017) as well as the investigation of empirical data (Ji et al. 2020; Grilli 2020; Descheemaeker & de Buyl 2020). It stands to reason that comparative population genetics could learn from such an approach. While our argument here is primarily statistical, a spiritually similar argument has been made regarding the use of effective models that coarse-grain over taxonomic details to identify quantitative patterns in microbial ecology and evolution (Good & Hallatschek 2018).

Ultimately it is necessary to take stock of the set of patterns that remain robust across phylogenetically diverged species within the human gut, allowing us to identify the evolutionary dynamics that universally occur. Here, we will briefly examine a few notable evolutionary patterns that have been observed across species.

##### i. Population Structure

The genetic composition of commensal bacteria varies considerably from host to host (Schloissnig et al. 2013; Truong et al. 2017; Costea et al. 2017), suggesting that bacteria do not rampantly migrate between hosts. Instead, for each species, hosts are typically colonized by a handful of strains that seem to be unique to each host (Garud et al. 2019; Schloissnig et al. 2013). The typical number of strains within a species is variable, likely reflecting the degree that strains can diverge and evolve sufficient ecological differences necessary to co-exist (Good et al. 2018). Though this within-host population structure does not seem to necessarily have bearing on across-host population structure, as the global biogeography of genetic variants can vary considerably across species. For example, the genetic diversity of *Eubacterium rectale* mirrors the genetic diversity of hosts (Truong et al. 2017; Tett et al. 2019; Nayfach et al. 2016; Costea et al. 2017; Karcher et al. 2020), whereas species such as *B. vulgatus* seem to show little or no geographic structure. The mechanisms responsible for variation in the global biogeography of species remains unclear. Vertical transmission from parents to infants may contribute, as strains from certain species are more likely to colonize and persist in infant guts, the genera *Bacteroides* and *Bifidobacterium* being noted examples (Lou et al. 2021). Though the benefit of being the first to colonize a host is likely temporary, as the majority of strains are replaced over several decades (Garud et al. 2019). Alternative mechanisms for varied levels of biogeography include traits that promote airborne transmission being restricted to certain lineages (Brown 2000) which may explain the variation in transmission rates among species (Brito et al. 2019), the interaction of the microbiome with the genetics of its host (Goodrich et al. 2016), and even the presence of spatial structure itself, which may promote the preservation of genetic variation (Pearce & Fisher 2019), though these hypotheses remain to be fully tested.

##### ii. Recombination

Although all bacteria reproduce clonally, the degree to which bacteria recombine varies widely. The recombination rate of a species determines whether populations evolve primarily via changes in genotype frequencies over time or as changes in the frequencies of individual alleles that are effectively independent (Neher & Shraiman 2011), which can have consequences for whether gene-specific versus genome-wide selective sweeps are more common (Shapiro et al. 2012; B. Jesse Shapiro 2016; Bendall et al. 2016). To quantify recombination in bacteria, researchers have begun to characterize the statistical association of alleles at different loci (i.e., linkage disequilibria), where the degree of association can be viewed a function of the recombination rate. For several species found in the gut, as well as environmental samples and pathogens, linkage disequilibria tends to decay as the genetic distance between a pair of loci increases, which suggests that recombination may be common (Crits-Christoph et al. 2020; Sakoparnig et al. 2021; Lin & Kussell 2019). Such rampant recombination suggests that while microbes reproduce clonally, the label “asexual” is a misnomer. Instead, microorganisms are increasingly being deemed as “quasi-sexual”, where a large number of loci evolve independently instead of as genotypes despite the clonal nature through which they are reproduced (Smith et al. 1993; Rosen et al. 2015; B Jesse Shapiro 2016). However, observed levels of linkage disequilibrium tend to be higher than what is expected under free recombination for many species (Garud et al. 2019). Some species may be truly clonal (Smith et al. 1993; Vos & Didelot 2009), while others are likely subject to additional evolutionary forces that can generate correlations between sites, such as demographic history and selection. Selection seems to play a particularly prominent role, where recent developments in population genetic theory provide the groundwork necessary for subsequent empirical investigation (Arnold et al. 2020; Good 2020). These forces will need to be disentangled to understand the full extent of recombination in the microbiome.

##### iii. Short term evolution within hosts

Recently, it was found that evolutionary changes can occur in the human gut microbiome on short timescales of just a few months and even days(Garud et al. 2019; Ghalayini et al. 2018; Roodgar et al. 2020; Zhao et al. 2019; Poyet et al. 2019; Yaffe & Relman 2020), and that strain replacements are generally rare over that timescale. These evolutionary changes modify the haplotypes of existing lineages and seem to derive from a mixture of *de novo* mutations and horizontal gene transfer via recombination. The recombination-seeded events are a unique mode of adaptation that highlight how a complex community can maintain a reservoir of adaptive genetic material, which may be particularly useful in rapidly fluctuating environments where evolution via *de novo* mutations may take a long time. Thus, complex communities may be able to modulate the mode and tempo of evolution of focal species (Madi et al. 2020). At longer time-scales, these evolutionary changes tend to give way to ecology, as strain replacements become common. So far, there does not seem to be any evidence that rates of evolution or strain replacement differ across species, though future work may identify any differences. Though these seemingly contrasting dynamics raise the broader point that being able to examine evolutionary dynamics across a range of timescales will ultimately require researchers to intuit what evolutionary and ecological dynamics are relevant on a given timescale and, ultimately, construct models that bridge separate dynamics.

##### iv. Purifying selection

By comparing haplotypes of a given species from different hosts, we can focus on patterns that are the outcome of evolutionary dynamics that have operated over an extended timescale. One such pattern is how the ratio of nonsynonymous to synonymous divergences (*d*_*N*_/*d*_*S*_) changes as synonymous divergence (*d*_*S*_) increases, which would indicate in what direction selection tends to dominate over an extended timescale. Looking at empirical data from the microbiome, it is clear that *d*_*N*_/*d*_*S*_ tends to decrease with increasing *d*_"_, suggesting that purifying selection tends to dominate as lineages diverge (Fig. 1a; Garud et al. 2019). Surprisingly, the shape of the relationship can be captured by an effective model of selection composed of two parameters: a single selection coefficient and the fraction of sites subject to selection (Garud et al. 2019). Some species clearly have values of *d*_*N*_/*d*_*S*_ that are further from this prediction than others, an observation that we will return to below. But as a first approximation we can say that purifying selection explains genome-wide patterns of genetic divergence across species within the gut (Fig. 1a).

**Fig. 1:**
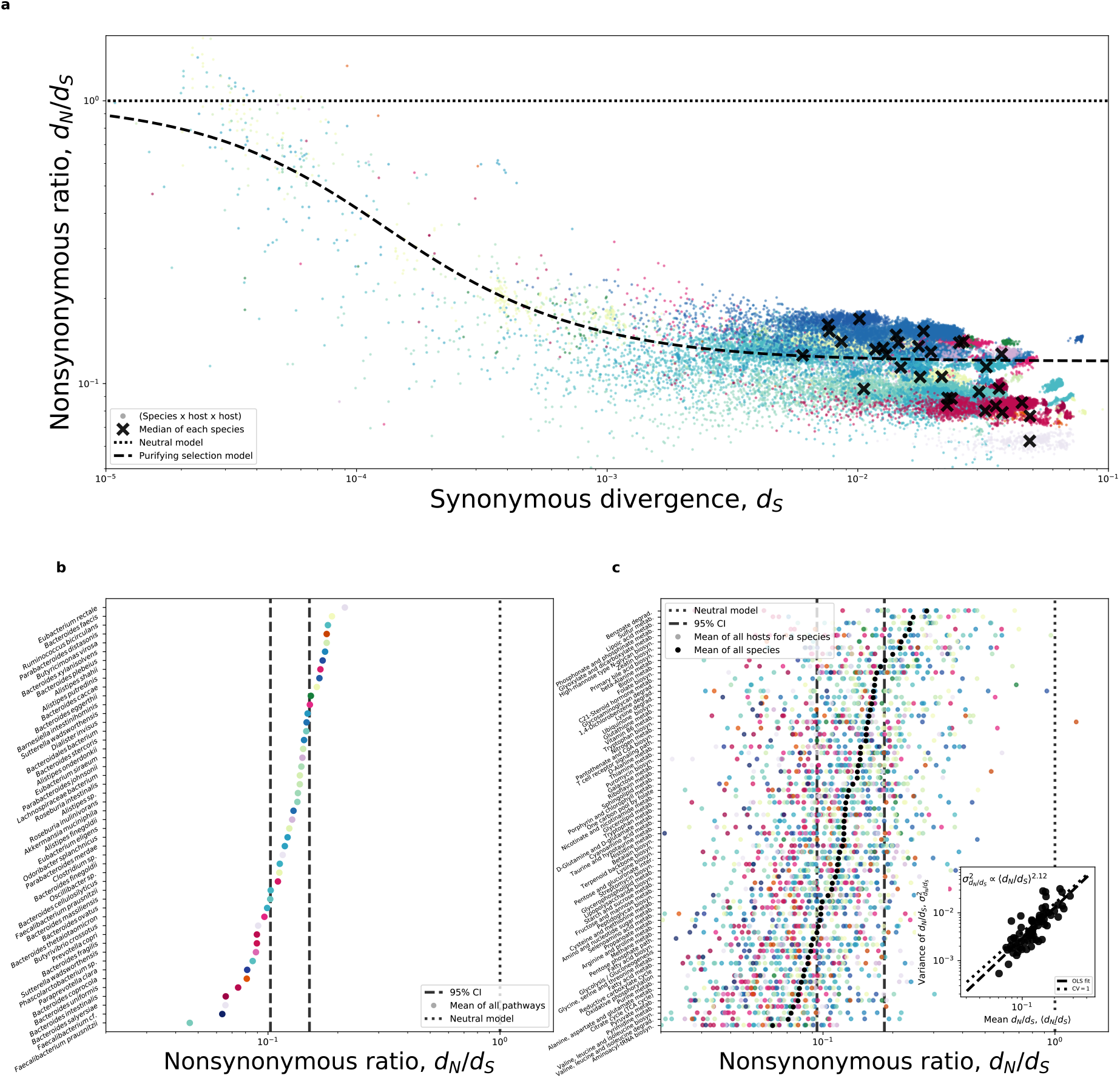
**a)** The relationship between synonymous divergence on the x-axis (*d*_*S*_) and the ratio of nonsynonymous and synonymous divergences (*d*_*N*_/*d*_*S*_) on the y-axis follows the form predicted by purifying selection across species (Eq. S8 from Garud, Good et al. (2019)). Though by color-coding individual species, we see that data points tend to be grouped by species identity, where certain species fall above or below the prediction. **b)** By grouping genes by their pathways and generating an appropriate null distribution via permutation, we can identify pathways that, under the assumptions of the model, are under stronger or weaker purifying selection than expected by chance. We can then examine how the mean *d*_*N*_/*d*_*S*_ (〈*d*_*N*_/*d*_*S*_〉) of a given pathway relates to its variance 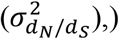, where the variance increases slightly faster than the square of 〈*d*_*N*_/*d*_*S*_〉, suggesting that the coefficient of variation is greater than one (inset figure in **b**). **c)** By inverting our permutation scheme, we can identify the set of species that are subject to stronger or weaker purifying selection than expected by chance.

#### 3) Identifying deviations from general trends

It may not be clear how a researcher can leverage universal evolutionary patterns to identify genes and species of interest. Indeed, interest in identifying exceptional species and targets of evolution is likely why many researchers compare species in the first place (Lawrie & Petrov 2014; Leffler et al. 2012; Huber et al. 2020). Here, we will re-examine the relationship between *d*_*S*_ and *d*_*N*_/*d*_*S*_ discussed earlier to illustrate how starting with a strong universal pattern can provide a backdrop against which we identify deviations from the overall trend. We see that certain species tend to fall above or below the prediction of an effective model of purifying selection (Fig. 1a). To identify species that are subject to stronger or weaker purifying selection than expected by chance, we first coarse-grain genes by their annotated metabolic pathways, providing a set of variables shared across species. We can then permute species-level observations within a given pathway and establish 95% confidence intervals, a non-parametric test that allows us to identify species with exceptional *d*_*N*_/*d*_*S*_ (Fig. 1b).

Though we coarse-grained genes by necessity, it allowed us to perform additional analyses in which we leverage information across multiple species. First, we can see that certain pathways typically have lower *d*_*N*_/*d*_*S*_ than others, suggesting that they are subject to stronger purifying selection (Fig. 1c). We can then identify pathways that are subject to stronger or weaker purifying selection than expected by chance by permuting values of *d*_*N*_/*d*_*S*_ across pathways within each species and establishing confidence intervals (see Supplemental Information). The results of this test align with our biological intuition, as essential pathways tend to be under stronger purifying selection (e.g., glycolysis, nucleotide biosynthesis, Krebs cycle, etc.), while pathways that rely on specific resources (e.g., sulfur metabolism) tend to be under relatively relaxed selection, an observation consistent with prior analyses using polymorphism data (Schloissnig et al. 2013).

Building off of our permutation analysis, because we have observations from many species, we can continue our comparative population genetic analyses and examine the statistical properties of *d*_*N*_/*d*_*S*_ across pathways. First, we can determine whether the relative spread of *d*_*N*_/*d*_*S*_ remains similar across pathways by examining whether the ratio of the standard deviation to the mean (i.e., the coefficient of variation) remains constant. We find that this is the case, as the mean *d*_*N*_/*d*_*S*_ of a pathway (〈*d*_*N*_/*d*_*S*_〉) across species is linear with respect to its variance (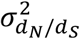; Fig. 1b). This observation is reminiscent of Taylor’s Law (Taylor 1961), a pattern often found in ecological systems (Grilli 2020). Similar to Taylor’s Law, we find that the slope of this relationship is not significantly different from two (*t* = 0.684; *P* = 0.248), which we can interpret as the mean being equal to the standard deviation across pathways 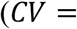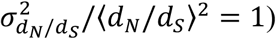. This observation suggests that the relative dispersion of *d*_*N*_/*d*_*S*_ remains constant as the overall level of constraint within a pathway is relaxed. Though values of *d*_*N*_/*d*_*S*_ across pathways are not independent, as the correlation in *d*_*N*_/*d*_*S*_ across pathways for a given pair of species tends to decay with phylogenetic distance (*β* = −0.104, *P* < 10^-6^), suggesting that the strength of purifying selection within a given essential pathway is moderately conserved through evolutionary time. However, this does not provide an explanation of why certain pathways are subject to stronger purifying selection than others, or why the strength of selection varies across lineages, a question that can likely only be answered by incorporating additional biological details about the pathways and species themselves (Bielawski & Yang 2004; Aguileta et al. 2009). Rather, it illustrates how investigating deviations from an empirical pattern can lead to novel findings.

### Future directions for comparative population genetics in the microbiome

The study of comparative population genetics in the human microbiome is nascent and full of potential. There are multiple avenues of progress in the microbiome field that will benefit comparative population genetics as a discipline. First, advances in sequencing technology will allow us to refine our estimates of important quantities. For example, long-read technologies, such as nanopore metagenomic sequencing, will allow researchers to quantify linkage between physically distant sites (Bharti & Grimm 2019; Zlitni et al. 2020; Yaffe & Relman 2020; Karst et al. 2021), providing higher resolution to uncover fundamental evolutionary processes of recombination, mutation, and adaptation within and across species. These advances, coupled with decreasing costs of library preparation (Baym et al. 2015), will allow researchers to sample large cohorts over time and observe how genotypes dissipate into alleles and reemerge via recombination over their sojourn times, enabling us to build more detailed evolutionary models.

Second, the fact that the gut exists as a physical environment is often overlooked. Environmental factors such as temperature (Groussin & Gouy 2011) as well as spatial structure (Tropini et al. 2017) can affect the evolutionary dynamics of microbial species. Even deceptively simple features of the gut such as its resemblance to a chemostat or the peristaltic mixing that arises due to digestion can produce complex ecological and evolutionary dynamics (Locey & Lennon 2019; Cremer et al. 2016).

Finally, there is arguably as much a need to examine the targets of molecular evolution as the general processes shaping genetic variation in the microbiome. The genes that contribute to adaptation ultimately encode physical aspects of cells, which means that subsequent experiments will be necessary to gain a more complete understanding on how adaptation in the gut proceeds and varies across species (Lynch & Trickovic 2020; Lynch et al. 2014). These advances and considerations will allow researchers to understand the general evolutionary dynamics of the microbiome, expanding the breadth and depth of comparative population genetics as a discipline

## Data Availability

The raw sequencing reads for the metagenomic samples used in this study were previously described (Garud, et al., 2019). The source code for figure generation and associated metadata is available on GitHub: https://github.com/garudlab/CompPopGenMicrobiomeReview.

## Acknowledgments

We thank Kirk Lohmueller for helpful discussions and John Connoly and Leah Briscoe for providing feedback on an earlier draft. This work was supported by the NSF Postdoctoral Research Fellowships in Biology Program under Grant No. 2010885.

## References

Advani M, Bunin G, Mehta P. 2018. Statistical physics of community ecology: a cavity solution to MacArthur’s consumer resource model. J. Stat. Mech. 2018:033406. doi: 10.1088/1742-5468/aab04e.

Aguileta G, Refrégier G, Yockteng R, Fournier E, Giraud T. 2009. Rapidly evolving genes in pathogens: methods for detecting positive selection and examples among fungi, bacteria, viruses and protists. Infect Genet Evol. 9:656–670. doi: 10.1016/j.meegid.2009.03.010.

Almeida A et al. 2019. A new genomic blueprint of the human gut microbiota. Nature. 568:499–504. doi: 10.1038/s41586-019-0965-1.

Arnold B et al. 2020. Fine-Scale Haplotype Structure Reveals Strong Signatures of Positive Selection in a Recombining Bacterial Pathogen. Mol Biol Evol. 37:417–428. doi: 10.1093/molbev/msz225.

Barbier M, Arnoldi J-F. 2017. The cavity method for community ecology. Ecology doi: 10.1101/147728.

Barreto HC, Sousa A, Gordo I. 2020. The Landscape of Adaptive Evolution of a Gut Commensal Bacteria in Aging Mice. Current Biology. 30:1102–1109.e5. doi: 10.1016/j.cub.2020.01.037.

Barroso-Batista J et al. 2020. Specific Eco-evolutionary Contexts in the Mouse Gut Reveal Escherichia coli Metabolic Versatility. Current Biology. 30:1049–1062.e7. doi: 10.1016/j.cub.2020.01.050.

Barroso-Batista J et al. 2014. The First Steps of Adaptation of Escherichia coli to the Gut Are Dominated by Soft Sweeps. PLoS Genetics. 10:e1004182. doi: 10.1371/journal.pgen.1004182.

Baym M et al. 2015. Inexpensive Multiplexed Library Preparation for Megabase-Sized Genomes Green, SJ, editor. PLoS ONE. 10:e0128036. doi: 10.1371/journal.pone.0128036.

Belkaid Y, Hand TW. 2014. Role of the microbiota in immunity and inflammation. Cell. 157:121–141. doi: 10.1016/j.cell.2014.03.011.

Bendall ML et al. 2016. Genome-wide selective sweeps and gene-specific sweeps in natural bacterial populations. ISME Journal. 10:1589–1601. doi: 10.1038/ismej.2015.241.

Bertels F, Leemann C, Metzner KJ, Regoes RR. 2019. Parallel Evolution of HIV-1 in a Long-Term Experiment. Molecular Biology and Evolution. 36:2400–2414. doi: 10.1093/molbev/msz155.

Bharti R, Grimm DG. 2019. Current challenges and best-practice protocols for microbiome analysis. Briefings in Bioinformatics. doi: 10.1093/bib/bbz155.

Bielawski JP, Yang Z. 2004. A Maximum Likelihood Method for Detecting Functional Divergence at Individual Codon Sites, with Application to Gene Family Evolution. J Mol Evol. 59:121–132. doi: 10.1007/s00239-004-2597-8.

Blount ZD, Borland CZ, Lenski RE. 2008. Historical contingency and the evolution of a key innovation in an experimental population of Escherichia coli. PNAS. 105:7899–7906. doi: 10.1073/pnas.0803151105.

Brito IL et al. 2019. Transmission of human-associated microbiota along family and social networks. doi: 10.1038/s41564-019-0409-6.

Brown LM. 2000. Helicobacter pylori: Epidemiology and routes of transmission. Epidemiologic Reviews. 22:283–297. doi: 10.1093/oxfordjournals.epirev.a018040.

Bruger EL, Marx CJ. 2018. A decade of genome sequencing has revolutionized studies of experimental evolution. Current Opinion in Microbiology. 45:149–155. doi: 10.1016/j.mib.2018.03.002.

Cain AK et al. 2020. A decade of advances in transposon-insertion sequencing. Nature Reviews Genetics. 21:526–540. doi: 10.1038/s41576-020-0244-x.

Clark AG et al. 2007. Evolution of genes and genomes on the Drosophila phylogeny. Nature. 450:203–218. doi: 10.1038/nature06341.

Costea PI et al. 2017. Subspecies in the global human gut microbiome. Molecular Systems Biology. 13:960. doi: 10.15252/msb.20177589.

Cremer J et al. 2016. Effect of flow and peristaltic mixing on bacterial growth in a gut-like channel. PNAS. 113:11414–11419. doi: 10.1073/pnas.1601306113.

Crits-Christoph A, Olm MR, Diamond S, Bouma-Gregson K, Banfield JF. 2020. Soil bacterial populations are shaped by recombination and gene-specific selection across a grassland meadow. The ISME Journal. 14:1834–1846. doi: 10.1038/s41396-020-0655-x.

Cvijović I, Good BH, Desai MM. 2018. The Effect of Strong Purifying Selection on Genetic Diversity. Genetics. 209:1235–1278. doi: 10.1534/genetics.118.301058.

Davydov EV et al. 2010. Identifying a High Fraction of the Human Genome to be under Selective Constraint Using GERP++. PLOS Computational Biology. 6:e1001025. doi: 10.1371/journal.pcbi.1001025.

Descheemaeker L, de Buyl S. 2020. Stochastic logistic models reproduce experimental time series of microbial communities Krishna, S & Walczak, AM, editors. eLife. 9:e55650. doi: 10.7554/eLife.55650.

D’Souza G et al. 2018. Ecology and evolution of metabolic cross-feeding interactions in bacteria. Nat. Prod. Rep. 35:455–488. doi: 10.1039/C8NP00009C.

Duvallet C, Gibbons SM, Gurry T, Irizarry RA, Alm EJ. 2017. Meta-analysis of gut microbiome studies identifies disease-specific and shared responses. Nature Communications. 8:1784. doi: 10.1038/s41467-017-01973-8.

Faith JJ et al. 2013. The long-term stability of the human gut microbiota. Science. 341. doi: 10.1126/science.1237439.

Fremin BJ, Bhatt AS. 2020. A combined RNA-Seq and comparative genomics approach identifies 1,085 candidate structured RNAs expressed in human microbiomes. bioRxiv. 2020.03.31.018887. doi: 10.1101/2020.03.31.018887.

Garud NR, Good BH, Hallatschek O, Pollard KS. 2019. Evolutionary dynamics of bacteria in the gut microbiome within and across hosts Gordo, I, editor. PLoS Biol. 17:e3000102. doi: 10.1371/journal.pbio.3000102.

Garud NR, Pollard KS. 2020. Population Genetics in the Human Microbiome. Trends in Genetics. 36:53–67. doi: 10.1016/j.tig.2019.10.010.

Gautam A et al. 2018. Altered fecal microbiota composition in all male aggressor-exposed rodent model simulating features of post-traumatic stress disorder. Journal of Neuroscience Research. 96:1311–1323. doi: 10.1002/jnr.24229.

Ghalayini M et al. 2018. Evolution of a dominant natural isolate of Escherichia coli in the human gut over the course of a year suggests a neutral evolution with reduced effective population size. Applied and Environmental Microbiology. 84. doi: 10.1128/AEM.02377-17.

Good BH. 2020. Linkage disequilibrium between rare mutations. bioRxiv. 2020.12.10.420042. doi: 10.1101/2020.12.10.420042.

Good BH, Hallatschek O. 2018. Effective models and the search for quantitative principles in microbial evolution. Current Opinion in Microbiology. 45:203–212. doi: 10.1016/j.mib.2018.11.005.

Good BH, Martis S, Hallatschek O. 2018. Adaptation limits ecological diversification and promotes ecological tinkering during the competition for substitutable resources. Proc Natl Acad Sci USA. 115:E10407–E10416. doi: 10.1073/pnas.1807530115.

Good BH, McDonald MJ, Barrick JE, Lenski RE, Desai MM. 2017. The dynamics of molecular evolution over 60,000 generations. Nature. 551:45–50. doi: 10.1038/nature24287.

Goodrich JK, Davenport ER, Waters JL, Clark AG, Ley RE. 2016. Cross-species comparisons of host genetic associations with the microbiome. doi: 10.1126/science.aad9379.

Gould SJ. 1990. Wonderful life: the Burgess Shale and the nature of history. Norton & Co: New York.

Goyal A, Bittleston LS, Leventhal GE, Lu L, Cordero OX. 2021. Interactions between strains govern the eco-evolutionary dynamics of microbial communities. bioRxiv. 2021.01.04.425224. doi: 10.1101/2021.01.04.425224.

Grilli J. 2020. Macroecological laws describe variation and diversity in microbial communities. Nature Communications. 11:4743. doi: 10.1038/s41467-020-18529-y.

Groussin M, Gouy M. 2011. Adaptation to Environmental Temperature Is a Major Determinant of Molecular Evolutionary Rates in Archaea. Molecular Biology and Evolution. 28:2661–2674. doi: 10.1093/molbev/msr098.

Hehemann JH et al. 2010. Transfer of carbohydrate-active enzymes from marine bacteria to Japanese gut microbiota. Nature. 464:908–912. doi: 10.1038/nature08937.

Herron MD, Doebeli M. 2013. Parallel Evolutionary Dynamics of Adaptive Diversification in Escherichia coli. PLoS Biology. 11. doi: 10.1371/journal.pbio.1001490.

Huber CD, Kim BY, Lohmueller KE. 2020. Population genetic models of GERP scores suggest pervasive turnover of constrained sites across mammalian evolution. PLOS Genetics. 16:e1008827. doi: 10.1371/journal.pgen.1008827.

Jakobsson HE et al. 2010. Short-Term Antibiotic Treatment Has Differing Long-Term Impacts on the Human Throat and Gut Microbiome. PLOS ONE. 5:e9836. doi: 10.1371/journal.pone.0009836.

Ji BW, Sheth RU, Dixit PD, Tchourine K, Vitkup D. 2020. Macroecological dynamics of gut microbiota. Nature Microbiology. 5:768–775. doi: 10.1038/s41564-020-0685-1.

Johnson ZI, Chisholm SW. 2004. Properties of overlapping genes are conserved across microbial genomes. Genome Res. 14:2268–2272. doi: 10.1101/gr.2433104.

Karcher N et al. 2020. Analysis of 1321 Eubacterium rectale genomes from metagenomes uncovers complex phylogeographic population structure and subspecies functional adaptations. Genome Biology. 21:138. doi: 10.1186/s13059-020-02042-y.

Karst SM et al. 2021. High-accuracy long-read amplicon sequences using unique molecular identifiers with Nanopore or PacBio sequencing. Nature Methods. 18:165–169. doi: 10.1038/s41592-020-01041-y.

Kenny DJ et al. 2020. Cholesterol Metabolism by Uncultured Human Gut Bacteria Influences Host Cholesterol Level. Cell Host and Microbe. 28:245–257.e6. doi: 10.1016/j.chom.2020.05.013.

Kimura M. 1983. The Neutral Theory of Molecular Evolution. 1st ed. Cambridge University Press doi: 10.1017/CBO9780511623486.

Korem T et al. 2015. Growth dynamics of gut microbiota in health and disease inferred from single metagenomic samples. Science. 349:1101–1106. doi: 10.1126/science.aac4812.

Lawrie DS, Petrov DA. 2014. Comparative population genomics: power and principles for the inference of functionality. Trends in Genetics. 30:133–139. doi: 10.1016/j.tig.2014.02.002.

Leffler EM et al. 2012. Revisiting an Old Riddle: What Determines Genetic Diversity Levels within Species? PLOS Biology. 10:e1001388. doi: 10.1371/journal.pbio.1001388.

Levins R, Lewontin RC. 1987. The dialectical biologist. Harvard University Press: Cambridge, Mass.

Levy SF et al. 2015. Quantitative evolutionary dynamics using high-resolution lineage tracking. Nature. 519:181–186. doi: 10.1038/nature14279.

Ley RE et al. 2005. Obesity alters gut microbial ecology. Proceedings of the National Academy of Sciences of the United States of America. 102:11070–11075. doi: 10.1073/pnas.0504978102.

Ley RE, Turnbaugh PJ, Klein S, Gordon JI. 2006. Microbial ecology: human gut microbes associated with obesity. Nature. 444:1022–1023. doi: 10.1038/4441022a.

Lieberman TD et al. 2014. Genetic variation of a bacterial pathogen within individuals with cystic fibrosis provides a record of selective pressures. Nature Genetics. 46:82–87. doi: 10.1038/ng.2848.

Lieberman TD. 2018. Seven Billion Microcosms: Evolution within Human Microbiomes. mSystems. 3:e00171–17, /msystems/3/2/msys.00171-17.atom. doi: 10.1128/mSystems.00171-17.

Lilja EE, Johnson DR. 2019. Substrate cross-feeding affects the speed and trajectory of molecular evolution within a synthetic microbial assemblage. BMC Evolutionary Biology. 19:129. doi: 10.1186/s12862-019-1458-4.

Lin M, Kussell E. 2017. Correlated Mutations and Homologous Recombination Within Bacterial Populations. Genetics. 205:891–917. doi: 10.1534/genetics.116.189621.

Lin M, Kussell E. 2019. Inferring bacterial recombination rates from large-scale sequencing datasets. Nat Methods. 16:199–204. doi: 10.1038/s41592-018-0293-7.

Lindblad-Toh K et al. 2011. A high-resolution map of human evolutionary constraint using 29 mammals. Nature. 478:476–482. doi: 10.1038/nature10530.

Locey KJ, Lennon JT. 2019. A Residence Time Theory for Biodiversity. The American Naturalist. 194:59–72. doi: 10.1086/703456.

Lou YC et al. 2021. Infant gut strain persistence is associated with maternal origin, phylogeny, and functional potential including surface adhesion and iron acquisition. bioRxiv. 2021.01.26.428340. doi: 10.1101/2021.01.26.428340.

Ludington WB, Ja WW. 2020. Drosophila as a model for the gut microbiome. PLOS Pathogens. 16:e1008398. doi: 10.1371/journal.ppat.1008398.

Lynch M et al. 2014. Evolutionary cell biology: Two origins, one objective. PNAS. 111:16990–16994. doi: 10.1073/pnas.1415861111.

Lynch M. 2007. The origins of genome architecture. Sinauer Associates: Sunderland, Mass.

Lynch M, Marinov GK. 2015. The bioenergetic costs of a gene. Proc Natl Acad Sci USA. 112:15690–15695. doi: 10.1073/pnas.1514974112.

Lynch M, Trickovic B. 2020. A Theoretical Framework for Evolutionary Cell Biology. Journal of Molecular Biology. 432:1861–1879. doi: 10.1016/j.jmb.2020.02.006.

Madi AN, Vos M, Murall CL, Legendre P. 2020. Title : Does diversity beget diversity in microbiomes ? bioRxiv. 1–26.

Martinez-Gutierrez CA, Aylward FO. 2019. Strong Purifying Selection Is Associated with Genome Streamlining in Epipelagic Marinimicrobia. Genome Biology and Evolution. 11:2887–2894. doi: 10.1093/gbe/evz201.

Milo R, Phillips R. 2016. Cell biology by the numbers. Garland Science, Taylor & Francis Group: New York, NY.

Moorjani P et al. 2016. A genetic method for dating ancient genomes provides a direct estimate of human generation interval in the last 45,000 years. PNAS. 113:5652–5657. doi: 10.1073/pnas.1514696113.

Nayfach S, Rodriguez-Mueller B, Garud N, Pollard KS. 2016. An integrated metagenomics pipeline for strain profiling reveals novel patterns of transmission and global biogeography of bacteria. bioRxiv. 031757. doi: 10.1101/031757.

Nayfach S, Shi ZJ, Seshadri R, Pollard KS, Kyrpides NC. 2019. New insights from uncultivated genomes of the global human gut microbiome. Nature. 568:505–510. doi: 10.1038/s41586-019-1058-x.

Neher RA, Shraiman BI. 2012. Fluctuations of Fitness Distributions and the Rate of Muller’s Ratchet. Genetics. 191:1283–1293. doi: 10.1534/genetics.112.141325.

Neher RA, Shraiman BI. 2011. Statistical genetics and evolution of quantitative traits. Rev. Mod. Phys. 83:1283–1300. doi: 10.1103/RevModPhys.83.1283.

Nguyen Ba AN et al. 2019. High-resolution lineage tracking reveals travelling wave of adaptation in laboratory yeast. Nature. 575:494–499. doi: 10.1038/s41586-019-1749-3.

Nicolaisen LE, Desai MM. 2012. Distortions in Genealogies Due to Purifying Selection. Molecular Biology and Evolution. 29:3589–3600. doi: 10.1093/molbev/mss170.

Pasolli E et al. 2019. Extensive Unexplored Human Microbiome Diversity Revealed by Over 150,000 Genomes from Metagenomes Spanning Age, Geography, and Lifestyle. Cell. 176:649–662.e20. doi: 10.1016/j.cell.2019.01.001.

Pathria RK, Beale PD. 2011. Statistical mechanics. 3rd ed. Elsevier/Academic Press: Amsterdam ; Boston.

Pearce MT, Fisher DS. 2019. Rapid adaptation in large populations with very rare sex: Scalings and spontaneous oscillations. Theoretical Population Biology. 129:18–40. doi: 10.1016/j.tpb.2017.11.005.

Pollard KS, Hubisz MJ, Rosenbloom KR, Siepel A. 2010. Detection of nonneutral substitution rates on mammalian phylogenies. Genome Res. 20:110–121. doi: 10.1101/gr.097857.109.

Powell JE, Leonard SP, Kwong WK, Engel P, Moran NA. 2016. Genome-wide screen identifies host colonization determinants in a bacterial gut symbiont. Proc Natl Acad Sci U S A. 113:13887–13892. doi: 10.1073/pnas.1610856113.

Poyet M et al. 2019. A library of human gut bacterial isolates paired with longitudinal multiomics data enables mechanistic microbiome research. Nat Med. 25:1442–1452. doi: 10.1038/s41591-019-0559-3.

Ramiro RS, Durão P, Bank C, Gordo I. 2020. Low mutational load and high mutation rate variation in gut commensal bacteria. PLOS Biology. 18:e3000617. doi: 10.1371/journal.pbio.3000617.

Romiguier J et al. 2014. Comparative population genomics in animals uncovers the determinants of genetic diversity. Nature. 515:261–263. doi: 10.1038/nature13685.

Roodgar M et al. 2020. Longitudinal linked read sequencing reveals ecological and evolutionary responses of a human gut microbiome during antibiotic treatment. bioRxiv. 2019.12.21.886093. doi: 10.1101/2019.12.21.886093.

Rosen MJ, Davison M, Bhaya D, Fisher DS. 2015. Fine-scale diversity and extensive recombination in a quasisexual bacterial population occupying a broad niche. Science. 348:1019–1023. doi: 10.1126/science.aaa4456.

Rowland I et al. 2018. Gut microbiota functions: metabolism of nutrients and other food components. Eur J Nutr. 57:1–24. doi: 10.1007/s00394-017-1445-8.

Ruiz L, Motherway MO, Lanigan N, Sinderen D van. 2013. Transposon Mutagenesis in Bifidobacterium breve: Construction and Characterization of a Tn5 Transposon Mutant Library for Bifidobacterium breve UCC2003. PLOS ONE. 8:e64699. doi: 10.1371/journal.pone.0064699.

Sakoparnig T, Field C, van Nimwegen E. 2021. Whole genome phylogenies reflect the distributions of recombination rates for many bacterial species Nourmohammad, A, editor. eLife. 10:e65366. doi: 10.7554/eLife.65366.

Sberro H et al. 2019. Large-Scale Analyses of Human Microbiomes Reveal Thousands of Small, Novel Genes. Cell. 178:1245–1259.e14. doi: 10.1016/j.cell.2019.07.016.

Scerri EML et al. 2018. Did Our Species Evolve in Subdivided Populations across Africa, and Why Does It Matter? Trends in Ecology & Evolution. 33:582–594. doi: 10.1016/j.tree.2018.05.005.

Schloissnig S et al. 2013. Genomic variation landscape of the human gut microbiome. Nature. 493:45–50. doi: 10.1038/nature11711.

Shapiro B Jesse. 2016. How clonal are bacteria over time? Evolutionary Biology doi: 10.1101/036780.

Shapiro B Jesse. 2016. How clonal are bacteria over time? Current Opinion in Microbiology. 31:116–123. doi: 10.1016/j.mib.2016.03.013.

Shapiro BJ et al. 2012. Population genomics of early events in the ecological differentiation of bacteria. Science. 335:48–51. doi: 10.1126/science.1218198.

Slobodkin LB. 1980. Growth and regulation of animal populations. 2d enl. ed. Dover Publications: New York.

Smith JM, Smith NH, O’Rourke M, Spratt BG. 1993. How clonal are bacteria? Proceedings of the National Academy of Sciences. 90:4384–4388. doi: 10.1073/pnas.90.10.4384.

Spanogiannopoulos P, Bess EN, Carmody RN, Turnbaugh PJ. 2016. The microbial pharmacists within us: A metagenomic view of xenobiotic metabolism. doi: 10.1038/nrmicro.2016.17.

Taylor LR. 1961. Aggregation, Variance and the Mean. Nature. 189:732–735. doi: 10.1038/189732a0.

Tenaillon O et al. 2016. Tempo and mode of genome evolution in a 50,000-generation experiment. Nature. 536:165–170. doi: 10.1038/nature18959.

Tett A et al. 2019. The Prevotella copri Complex Comprises Four Distinct Clades Underrepresented in Westernized Populations. Cell Host and Microbe. 26:666–679.e7. doi: 10.1016/j.chom.2019.08.018.

Thibault D et al. 2019. Droplet Tn-Seq combines microfluidics with Tn-Seq for identifying complex single-cell phenotypes. Nature Communications. 10:5729. doi: 10.1038/s41467-019-13719-9.

Thomas AM et al. 2019. Metagenomic analysis of colorectal cancer datasets identifies cross-cohort microbial diagnostic signatures and a link with choline degradation. Nature Medicine. 25:667–678. doi: 10.1038/s41591-019-0405-7.

Tropini C, Earle KA, Huang KC, Sonnenburg JL. 2017. The Gut Microbiome: Connecting Spatial Organization to Function. Cell Host and Microbe. 21:433–442. doi: 10.1016/j.chom.2017.03.010.

Truong DT, Tett A, Pasolli E, Huttenhower C, Segata N. 2017. Microbial strain-level population structure and genetic diversity from metagenomes. Genome Res. 27:626–638. doi: 10.1101/gr.216242.116.

Vallianou NG, Stratigou T, Tsagarakis S. 2018. Microbiome and diabetes: Where are we now? Diabetes Res Clin Pract. 146:111–118. doi: 10.1016/j.diabres.2018.10.008.

Vos M, Didelot X. 2009. A comparison of homologous recombination rates in bacteria and archaea. ISME Journal. 3:199–208. doi: 10.1038/ismej.2008.93.

Xue KS et al. 2017. Parallel evolution of influenza across multiple spatiotemporal scales Neher, RA, editor. eLife. 6:e26875. doi: 10.7554/eLife.26875.

Yaffe E, Relman DA. 2020. Tracking microbial evolution in the human gut using Hi-C reveals extensive horizontal gene transfer, persistence and adaptation. Nat Microbiol. 5:343–353. doi: 10.1038/s41564-019-0625-0.

Yano JM et al. 2015. Indigenous Bacteria from the Gut Microbiota Regulate Host Serotonin Biosynthesis. Cell. 161:264–276. doi: 10.1016/j.cell.2015.02.047.

Zhao S et al. 2019. Adaptive Evolution within Gut Microbiomes of Healthy People. Cell Host & Microbe. 25:656–667.e8. doi: 10.1016/j.chom.2019.03.007.

Zheng W et al. 2020. Microbe-seq: high-throughput, single-microbe genomics with strain resolution, applied to a human gut microbiome. bioRxiv. 2020.12.14.422699. doi: 10.1101/2020.12.14.422699.

Zhu B, Wang X, Li L. 2010. Human gut microbiome: The second genome of human body. doi: 10.1007/s13238-010-0093-z.

Zhu Q et al. 2019. Phylogenomics of 10,575 genomes reveals evolutionary proximity between domains Bacteria and Archaea. Nat Commun. 10:5477. doi: 10.1038/s41467-019-13443-4.

Zimmermann M, Zimmermann-Kogadeeva M, Wegmann R, Goodman AL. 2019. Separating host and microbiome contributions to drug pharmacokinetics and toxicity. Science. 363. doi: 10.1126/science.aat9931.

Zlitni S et al. 2020. Strain-resolved microbiome sequencing reveals mobile elements that drive bacterial competition on a clinical timescale. Genome Medicine. 12:50. doi: 10.1186/s13073-020-00747-0.

