## Supplemental Information for "Comparative Population Genetics in the Human Gut Microbiome"

### Calculating synonymous and nonsynonymous divergence

Synonymous and nonsynonymous divergences were calculated using a previously published approach and pipeline (Garud et al. 2019). Values of $d_N$ and $d_S$ at the metabolic level were calculated using code from the same pipeline, where KEGG annotations were obtained from PATRIC (Davis et al. 2020). KEGG genes were mapped to their higher-level metabolic pathways and $d_N$ and $d_S$ estimates for each pathway was corrected by the sum of gene sizes (i.e., target size for mutations).

### Examining how correlation in $d_N/d_S$ decays with phylogenetic distance.

The 16S rRNA sequence of each species was obtained from PATRIC (Davis et al., 2020). All sequences were aligned using MUSCLE v3.8.1551 (Edgar 2004) and aligned sites with >90% empty values were removed. Phylogenetic reconstruction was performed using Randomized Axelerated Maximum Likelihood v8.2.11 (RAxML) (Stamatakis 2014) with the General Time

Reversible model, gamma distributed rate variation, and bootstrap convergence criteria set to autoMRE. The 16S rRNA sequence of *Prochlorococcus marinus* subsp. *marinus* str. CCMP1375 (NCBI accession number NC005042) was used as an outgroup. Phylogenetic distance was calculated using the ETE Toolkit (Huerta-Cepas et al. 2016). Significance of the slope of the distance-decay relationship was established by comparing the slope inferred via ordinary least squares regression to a distribution of null slopes generated by randomly permuting species-pair labels.

### References

Davis JJ et al. 2020. The PATRIC Bioinformatics Resource Center: expanding data and analysis capabilities. Nucleic Acids Research. 48:D606–D612. doi: 10.1093/nar/gkz943.

Edgar RC. 2004. MUSCLE: multiple sequence alignment with high accuracy and high throughput. Nucleic Acids Res. 32:1792–1797. doi: 10.1093/nar/gkh340.

Garud NR, Good BH, Hallatschek O, Pollard KS. 2019. Evolutionary dynamics of bacteria in the gut microbiome within and across hosts Gordo, I, editor. PLoS Biol. 17:e3000102. doi: 10.1371/journal.pbio.3000102.

Huerta-Cepas J, Serra F, Bork P. 2016. ETE 3: Reconstruction, Analysis, and Visualization of Phylogenomic Data. Molecular Biology and Evolution. 33:1635–1638. doi: 10.1093/molbev/msw046.

Stamatakis A. 2014. RAxML version 8: a tool for phylogenetic analysis and post-analysis of large phylogenies. Bioinformatics. 30:1312–1313. doi: 10.1093/bioinformatics/btu033.

### Figures

**Fig. S1:** The correlation of nonsynonymous ratios across pathways tends to decay with phylogenetic distance for a given pair of species.

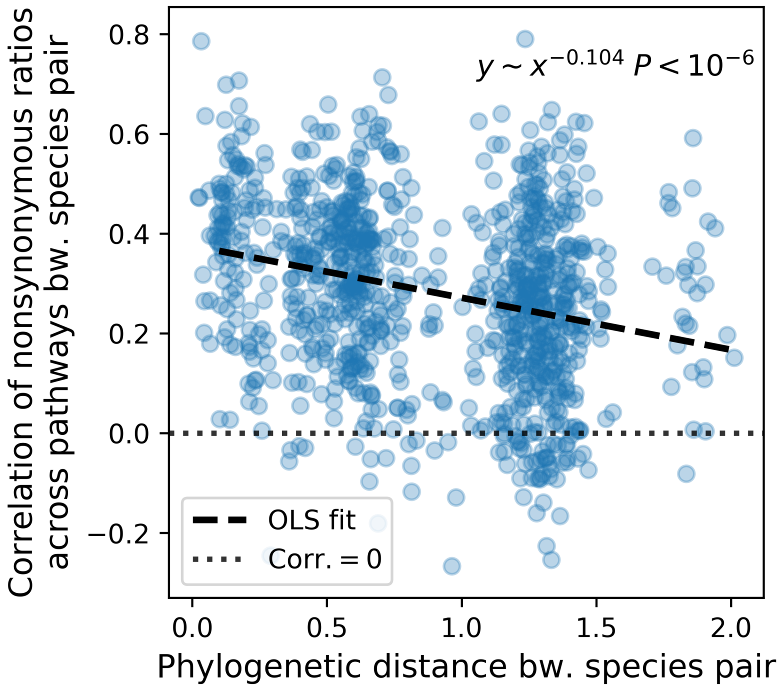
